# Genomic evidence supports a clonal diaspora model for metastases of esophageal adenocarcinoma

**DOI:** 10.1101/454306

**Authors:** Ayesha Noorani, Martin Goddard, Jason Crawte, Ludmil B. Alexandrov, Xiaodun Li, Maria Secrier, Matthew D. Eldridge, Lawrence Bower, Jamie Weaver, Pierre Lao-Sirieix, Inigo Martincorena, Irene Debiram-Beecham, Nicola Grehan, Shona MacRae, Shalini Malhotra, Ahmad Miremadi, Tabitha Thomas, Sarah Galbraith, Lorraine Petersen, Stephen D. Preston, David Gilligan, Andrew Hindmarsh, Richard H. Hardwick, OCCAMS Consortium, Michael R. Stratton, David C. Wedge, Rebecca C. Fitzgerald

**Affiliations:** MRC Cancer Unit, University of Cambridge, Biomedical Campus, Cambridge, CB2 OXZ, UK; Department of Histopathology, Papworth Hospital NHS Trust, Cambridge, CB23 3RE, UK; Theoretical Biology and Biophysics (T-6), Los Alamos National Laboratory, New Mexico, 87545, USA; Cancer Research UK Cambridge Research Institute, Cambridge, CB2 0RE, UK; Wellcome Trust Sanger Institute, Cambridge, CB10 1SA, UK; Department of Histopathology, Cambridge University Hospitals NHS Foundation Trust, Cambridge, CB2 0QQ, UK; Arthur Rank Hospice Charity, Cambridge, CB22 3FB, UK; Department of Palliative Care, Cambridge University Hospitals NHS Foundation Trust, Cambridge, CB2 0QQ, UK; Oncology Centre, Cambridge University Hospitals NHS Foundation Trust, Cambridge, CB2 0QQ, UK; Oesophago-Gastric Centre, Cambridge University Hospitals NHS Foundation Trust, Cambridge, CB2 0QQ, UK; Big Data Institute, University of Oxford, Oxford, OX3 7LF, UK; Oxford NIHR Biomedical Research Centre, Oxford, OX4 2PG, UK

## Abstract

Continual evolution of cancer makes it challenging to predict clinical outcomes. Highly varied and unpredictable patient outcomes in esophageal adenocarcinoma (EAC) prompted us to question the pattern and timing of metastatic spread. Whole genome sequencing and phylogenetic analysis of 396 samples across 18 EAC cases demonstrated a stellate pattern on the phylogenetic trees in 90% cases. The age-dependent trinucleotide signature, which can serve as a molecular clock, was absent or reduced in the stellate branches beyond the trunk in most cases (p<0.0001). Clustering of lymph nodes and distant metastases (n=250) demonstrated samples sharing a common clonal origin were widely dispersed anatomically. Metastatic subclones at autopsy were present in tissue and blood samples from earlier time-points. We infer that metastasis occurs rapidly across multiple sites, constituting a model of metastatic spread we term clonal diaspora. This has implications for understanding metastatic progression, clinical staging and patient management.

## Introduction

In cancer, metastatic spread to distant sites accounts for the majority of deaths (Sporn, 1996). Understanding the anatomical extent of disease is essential to determine the optimum treatment strategy for any given patient. This is difficult in practice since cancer continually evolves at a microscopic scale, often beyond the resolution of clinical imaging techniques. Furthermore, the patterns of metastatic spread are often unpredictable in terms of time-course and anatomical location. Treatments may therefore be unnecessarily toxic (e.g. radical lymphadenectomy and high dose chemotherapy) or lead to under-treatment with high recurrence rates (Lou et al., 2013; Matsuda et al., 2017; Waterman et al., 2004).

Esophageal cancer is the sixth most common cause of cancer-related death worldwide and the current median survival time is still <1 year despite advances in treatment (Smyth et al., 2017). Incidence rates for esophageal adenocarcinoma (EAC) have risen sharply and it is now the predominant form in developed countries. Prognosis is highly variable for EAC patients and with a wide range in the proportion of patients surviving beyond 5 years (18-47% in patients with lymph node involvement) making it difficult to advise patients when embarking on a long course of grueling treatment (Cunningham et al., 2008; Waterman et al., 2004). Despite the clinical classification of EAC as curative or non-curative, depending on the location of associated lymph nodes (Japanese Gastric Cancer, 2011) and involvement of solid organs, controversy exists in the field concerning whether radical lymph node dissection improves outcome (Matsuda et al., 2017; Stiles et al., 2012; Waterman et al., 2004).

Intratumor heterogeneity has been widely reported in human cancer with the first formal description of clonal evolution espoused by Peter Nowell in 1976 (Nowell, 1976). Theories of tumor evolution attempt to understand how tumor cell populations respond to selective pressures (Greaves and Maley, 2012). Subsequently there has been much debate about the models of tumor evolution, including linear, branching, neutral and punctuated evolution (Davis et al., 2017; Klein, 2009). With the advent of genome-wide sequencing methods recent large-scale efforts have been made to delineate different models of evolution, summarized in Table S1. Knowledge of how genetic diversity emerges over time as metastases develop remains limited, in part due to the challenge in collecting multiple samples over space and time from cancer patients.

To understand the evolution of EAC, we designed a prospective study with extensive sampling over-time including samples from diagnosis, surgery for operative cases and warm autopsy. We used whole genome sequencing at high (50x) and shallow (1x) coverage to interrogate the clonal architecture across time and space. The overall study design, sampling and sequencing strategy are shown in Figure S1.

## Results

### Genomic architecture of 18 cases

Eighteen cases were included and the clinical demographics of these cases are shown in Table S2 and S3, with details of the individual samples used given in Table S4 and S5. In the first part of the study (Step 1, Figure S1C) we used 50x WGS to construct a phylogenetic tree for each case to understand the relationship between the primary and metastases (Figure 1 and Figures S2 and S3). Mutation clustering was performed, and the fractions of tumor cells carrying each set of mutations (Cancer Cell Fraction, CCF) within each sample were used to determine: 1) the clonal and sub-clonal architecture of each tumor (subclonal CCF <95%, clonal CCF ≥ 95%); 2) the hierarchy of events; and 3) the distance of these sub-clonal or number of mutations on each branch of the tree (Figure S1C). The CCF of each clone and subclone is shown in Tables S6 and S7 along with the total number of single nucleotide variants (SNVs) and the tumor purity estimated using the Battenberg algorithm in Table S8. The confidence intervals of the CCFs these clones and subclones are shown in Supplementary Tables S9. All tissue samples undergoing WGS (all snap frozen except diagnostic biopsies which were FFPE archival samples) were also dually scored by expert esophageal histopathologists using the standards set by the International Cancer Genome Consortium (ICGC). Further macro dissection was performed for low cellularity samples to achieve a cellularity >70% to avoid bias (Supplementary Methods).

**Figure 1.**
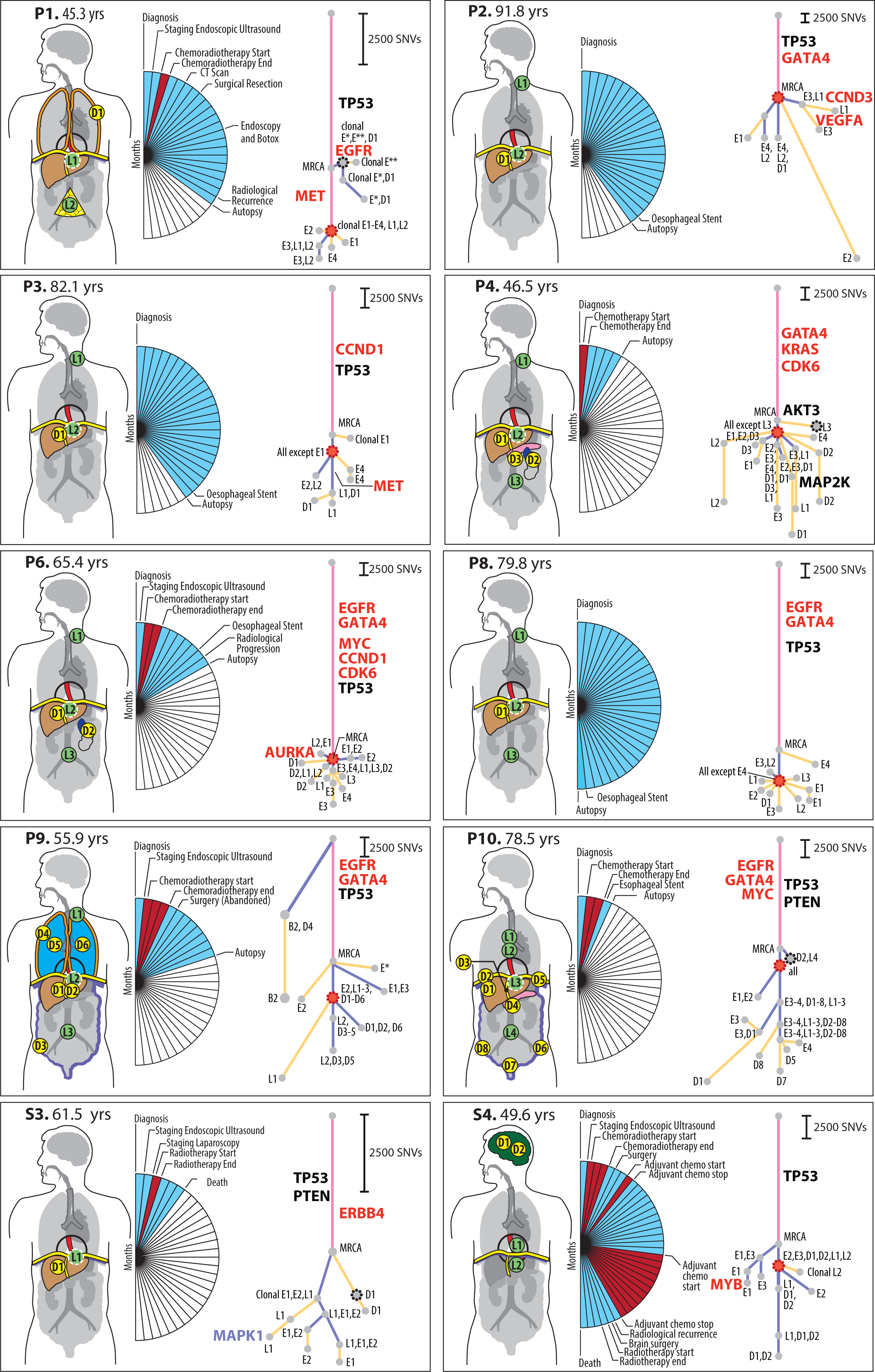
Phylogenetic Analysis of ten cases with nodal and distant metastases. Patient body maps (S for surgical case and P for rapid autopsy) are shown. Green circles denote lymph node metastases and yellow circles distant metastases. The labels within each circle describe the specific location (see Table S4 and S5 for precise anatomical descriptions). An organ is shown in color if metastases were sequenced from that case. The adjacent wedged semi-circle depicts the clinical timelines for each patient. Each wedge corresponds to one month; blue wedges indicate the total lifetime of the patient and red wedges indicate periods of therapy. Phylogenetic trees for each patient are shown and details of how these trees were constructed are provided in Supplementary methods and Supplementary Figures S12A and B; pink = truncal events shared by all samples, purple = branch events shared by more than one sample, yellow = leaves, events unique to a sample. The circle at the end of a trunk, branch or leaf represents a clone or subclone. Each clone or subclone is annotated to show which samples it is present in, where E1-E4 are samples from the primary esophageal tumor, L1-L4 are lymph nodes, and D1-8 are distant metastases - the numbering corresponds to the adjacent body map. A subclone annotated with E1, L2 for example indicates that this subclone is seen only in samples E1 and L2. The precise CCF of each subclone and clone (barring the MRCA) is shown in supplementary Tables S6 and S7. The length of the branches of the tree are reflective of the number of SNVs in the subclone/clone. The scales adjacent to each case are relative, given the variable number of SNVs per case. Trees are annotated with potential driver events, black: missense variants, red: amplifications. Gray dots outlined with a black dashed line denote the first subclone/clone to metastasize that would be classified as non-curative based on anatomical location. Red dots mark the stellate pattern on the phylogenetic tree.

These analyses enabled us to construct phylogenetic trees, in which the trunk represents the mutations common to all samples with length proportional to the number of mutations required for malignant transformation in that case. Branches represent subclones whose mutations are not found in every cancer cell. In all cases we observed a long trunk compared to the rest of the tree (median 19,034 SNVs, IQR 11,299-63,908), consistent with previous studies in EAC (Gerstung et al., 2017; Murugaesu et al., 2015). The median size of clusters across all cases was 3,069 SNVs (IQR1332-63908) and only 2/157 clusters contained fewer than 200 SNVs (case P5 and case S1, Figure S2).

The key driver events (Dulak et al., 2012; Secrier et al., 2016) are depicted on each phylogenetic tree (Figure 1 and Figure S2). The most frequent events that have previously been classed as drivers (Dulak et al., 2012; Frankell et al., 2018; Secrier et al., 2016) occurred in the trunks of the phylogenetic trees. TP53 was mutated in the trunk of 16 out of 18 cases, consistent with our knowledge of the disease (Dulak et al., 2013; Nones et al., 2014; Ross-Innes et al., 2015; Secrier et al., 2016; Weaver et al., 2014). Amplifications (gene names in red) were often truncal, but were also observed on the branches of the phylogenetic tree, providing evidence of divergence further down the evolutionary lineage (Figure 1, Figure S2). The majority of events in driver genes were copy number alterations rather than missense variants (Figure 1, Figure S2) (Frankell et al., 2018; Nones et al., 2014; Secrier et al., 2016). There was no significant difference in the overall number of structural variants between primary and metastatic samples (p=0.41, generalized linear model), (Figure S4b). However, a larger proportion of structural variants in metastatic samples were retro-transpositions of mobile elements compared with those in the primary samples (p=0.045, Figure S4c). This contrasts with pancreatic cancer, where deletions and fold-back inversions are more common, and breast cancer where tandem duplications dominate (Yates et al., 2015). Furthermore, the proportion of SVs found uniquely in metastases or in primary sites was higher than that of SNVs (Figure 1, Figure S4a), suggesting an increase in genomic instability in later stages of the disease. However, it cannot be ruled out that some SVs have not been identified in every sample as a result of lower sensitivity in the detection of SVs than SNVs.

Across the eighteen cases, 8 mutational signatures were observed, with varying prevalence and consistent with previous studies (Figure S5 (Ajani et al., 2015; Mariette et al., 2003; Sottoriva et al., 2013; Yachida et al., 2010). None of the signatures observed have been associated with treatment with alkylating antineoplastic agents (Alexandrov et al., 2013), platinum therapy (Liu et al., 2017) or radiation therapy (Behjati et al., 2016).

### Sub-cohort analysis of cases with local and distant spread

Ten of eighteen patients (S3, S4, P1-4, P6, P8-10) had nodal and solid organ metastases, allowing a direct comparison of the genomic architecture between different metastatic sites (Figure 1).

In four of these ten cases, an isolated clone or subclone confined to distant metastases shared the highest congruence to the most recent common ancestor (MRCA), depicted as a dashed black node on the first branch of the phylogenetic tree (P1, P4, P10, S3 in Figure 1). In P1, this subclone was shared between the primary tumor and a pleural metastasis. In S3 and P4, the clone involved in this isolated seeding was identified at a single distant site and not in the primary tumor (S3: liver metastasis (D1), P4: para-aortic lymph node (L3)); both events would render a patient’s management palliative. Interestingly, in P9 a subclone was found in a premalignant area of Barrett’s esophagus and a pleural metastasis but not in any of four areas of the primary tumor subject to 50X WGS. This lineage shares no variants with the main lineage and appears to be an independent second cancer. While undetected in any of the adenocarcinoma samples, it is plausible that this arose from an unsampled area of the primary tumor (Figure 1). The high CCF (>0.9) in both these cases suggests that these mutations developed *de novo* in metastatic sites soon after dissemination. In P10, the early seeding cluster was shared between a distant para-aortic node and a sub-clonal metastasis in the right hemi-diaphragm. This isolated seeding event showed little divergence from the MRCA (median 1913 SNVs, IQR 1540-1421) and suggests early seeding to distant metastases.

A striking observation was that 9/10 cases had a clone (outlined in red on the phylogenetic tree) that was followed by a dispersion of multiple subclones from the primary to discrete metastatic sites in a stellate pattern on the phylogenetic tree. The subclones forming this distinct pattern were located in both primary and metastatic tissue in eight cases (P1, P2, P3, S4, P4, P6, P8, P10) and in P9 were unique to metastases (Figure 1). P9 harbored sub-clonal CCFs in multiple sites, which could also indicate metastasis-to-metastasis seeding and further evidence for this was sought in the following parts of our study. The only case lacking a stellate pattern on the phylogenetic tree was S3, a non-autopsy case with limited tissue sampling. The early distant seeding in S3 is consistent with a pattern of parallel evolution (Figure 1).

### Shallow whole genome sequencing to assess spatial spread of EAC

In the second step of the study we aimed to elucidate the relative timing of metastatic events and to do this we performed 1X WGS in a further 250 tissue samples from 6 autopsy cases (Figure 2, Figure S1B). We did not call new mutations, as this would not be possible at 1X sequencing, but used this method to detect the spread of clones and subclones previously identified using 50X WGS (validation of methods in Figures S6 and S7). The samples used for this part of the study are outlined in Table S10. The median size of clusters (identified at 50X WGS) that we aimed to detect using 1X WGS was 3,784 (IQR 1966-49955). Sample sites were grouped according to their similarity based on the presence of subclones and clones previously detected with 50X WGS (Supplementary Methods, Shallow Whole Genome Sequencing for Subclone Identification). The resulting groups of samples are color coded and numbered, and the distribution of sample sites in these groups is shown on the adjacent body map. (Figure 2, see also Supplementary Methods). The most striking observation is that samples that grouped together based on shared clonal origins were widely dispersed anatomically.

**Figure 2.**
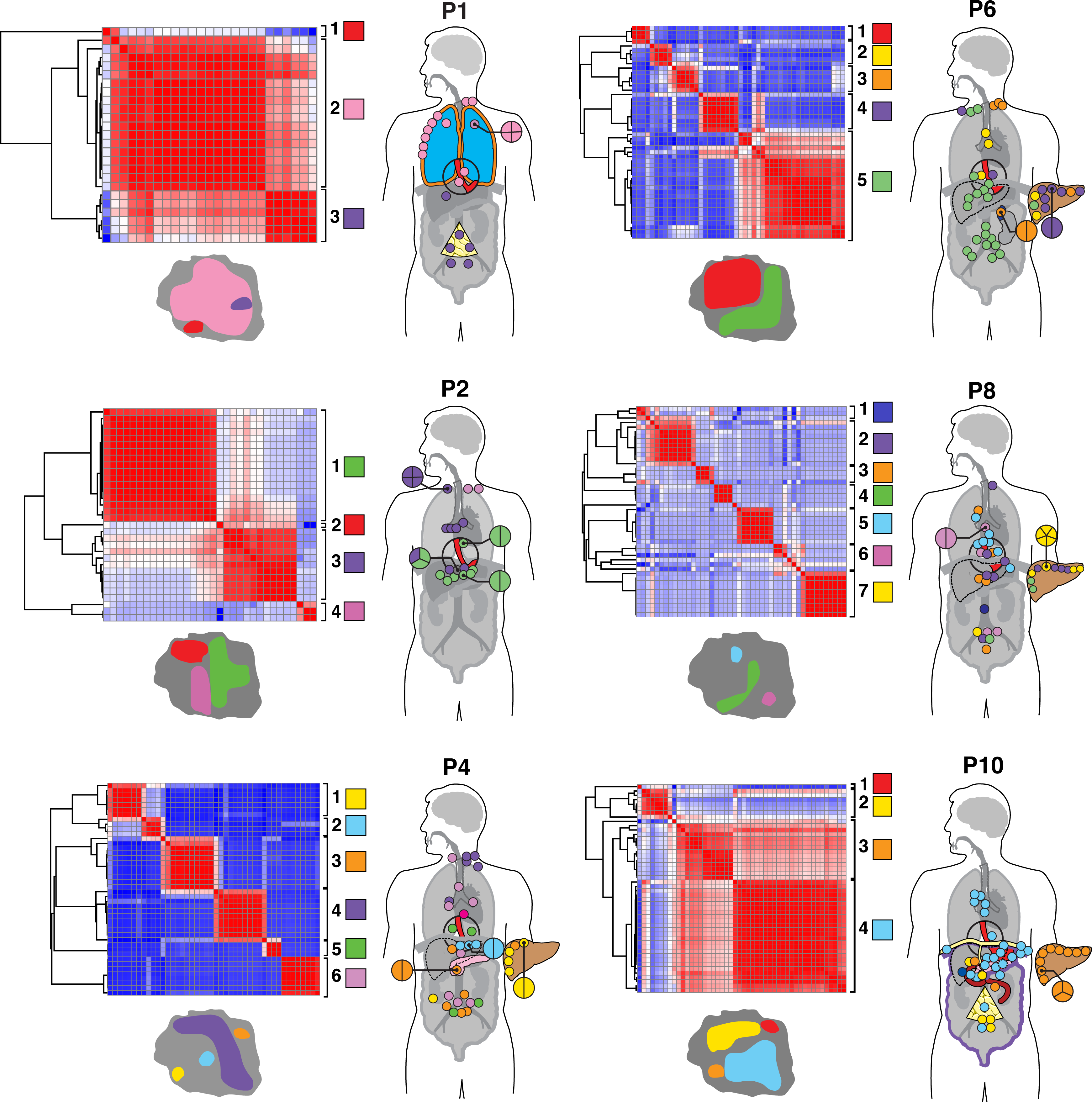
1X WGS and similarity matrix clustering of 250 further tissue samples from six cases. 1X WGS was performed at an average depth of 1x to track subclones and clones previously discovered using 50X WGS. Pearson correlation similarity matrix clustering was performed on all samples for each case (plotted against each other) with red indicating sample similarity (r=1) and blue indicating dissimilarity (r=−1). Sample sites used in this part of the sequenced. For example, liver metastases were only seen in P4,P6,P8,P10. Similarly, P2 had lymph nodes only (only colored dots are seen which represent lymph nodes, no solid organs are highlighted). Clustering was performed based on the presence of subclones and clones already detected using 50X WGS and distinct clusters were identified for each case as demonstrated by the adjacent key per case (each group is both colored and numbered). Samples are displayed on the adjoining body maps for which the color coding corresponds to the genomic clustering in the adjacent heatmap. Sites with multiple samples are magnified and the division of samples shown. Maps of the primary tumor with representation of metastatic subclones are shown with each case, with the colors of the subclones being the same as those in the matrix and body map. Areas shaded red in the primary tumor represent subclones that were not detected in the metastatic samples that underwent 1X WGS and were instead confined to areas of the primary tumor.

Four out of six cases with extensive spatial sampling (Figure 2) had liver metastases evaluated and 3 of these contained samples that were more similar to local lymph node metastases than neighboring liver metastases (P4, P6, P8 but not P10). The high number of groups within the liver (up to 4 in P6) suggested seeding by multiple subclones (seen in P4, P6, P8), whereas the single group (orange, number 3) in the liver of P10 indicated a single clonal expansion.

A comparison of lymph node location and genomic contiguity revealed no evidence of tropism, i.e. genomically similar lymph nodes did not occupy nearby anatomical locations. Lymph nodes above and below the diaphragm were frequently seeded from common events (P2: clusters 1, 3; P4: clusters 5, 6; P6: cluster 5; P8: clusters 2, 3, 6; P10: cluster 4), at odds with a progression from local to distant nodes. Similarly, a comparison of lymph node and solid organ metastases revealed scant evidence for tropism, with the exception of P1 (Supplementary Methods). In this cancer, separate subclones seeded lymph node and pleural metastases (Figures 1 and 2). However, the distant metastasis (D1) was seen to branch earlier than the lymph node metastasis in the evolutionary tree in our 50X WGS analysis (Figure 1).

We further traced regions of the primary tumor at autopsy with similar subclonal compositions to each of the groups of metastases, shown as adjacent tumor maps (Figure 2, bottom left of each case). Subclones occupied discrete, spatially distinct areas in the primary tumor.

### Timing of metastatic spread

To examine the timing of metastatic spread we analyzed the mutational signatures as well as comparing the mutations present at autopsy with those observed in the diagnostic biopsy samples and longitudinally collected plasma samples (Figure S1C).

Signature 1 arises from the enzymatic deamination of methylated cytosines which is an endogenous process that occurs continuously in both healthy and cancerous cells. This has been shown to act as a molecular clock, (Alexandrov et al., 2015; Alexandrov et al., 2013; Blokzijl et al., 2016; Gao et al., 2016; Letouze et al., 2017; Lodato et al., 2018), and was therefore used here as a method to examine the temporal relationship between metastases. Using a previously described method for deconvolving mutational signatures (Alexandrov et al., 2015), we observed that signature 1 was present in the trunk but absent in all subclones that constituted the stellate pattern on the phylogenetic tree (following the red clone in Figure 1) for P2, P4, P6, P9, P10, S4 and it was significantly reduced for P1 (21% to 3%) and P3 (16% to 9%) (Wilcoxon signed rank test p=0.039, Figure S15). Suspecting that the number of signature 1 mutations in branch subclones was below the resolution of our deconvolution methods, we identified the number of mutations with the characteristic feature of signature 1, i.e. C>T mutations at a CpG context, along the trunk to the stellate pattern and on the longest branch leading from the stellate pattern. With the exception of P8, the proportion of mutations with this feature was significantly lower post stellate pattern (p <9.1e-5, Chi-squared test) and the median proportion of such mutations occurring prior to the stellate pattern was 0.911 (Figure 3B). Thus, in the majority of cases one might deduce that very little time has elapsed between the appearance of the cell that is ancestral to disseminating cells and the individual cells that seeded each of the metastases. The substantial number of mutations arising from other mutational processes later in the evolutionary history (Table S11) suggests an increase in the activity of other processes. Of note, there was an increase in the proportion of signature 3 in subclonal SNVs compared to clonal SNVs, with this signature being associated with the failure of DNA double strand break repair (Wilcoxon signed rank test p=0.019, Figure S15).

**Figure 3.**
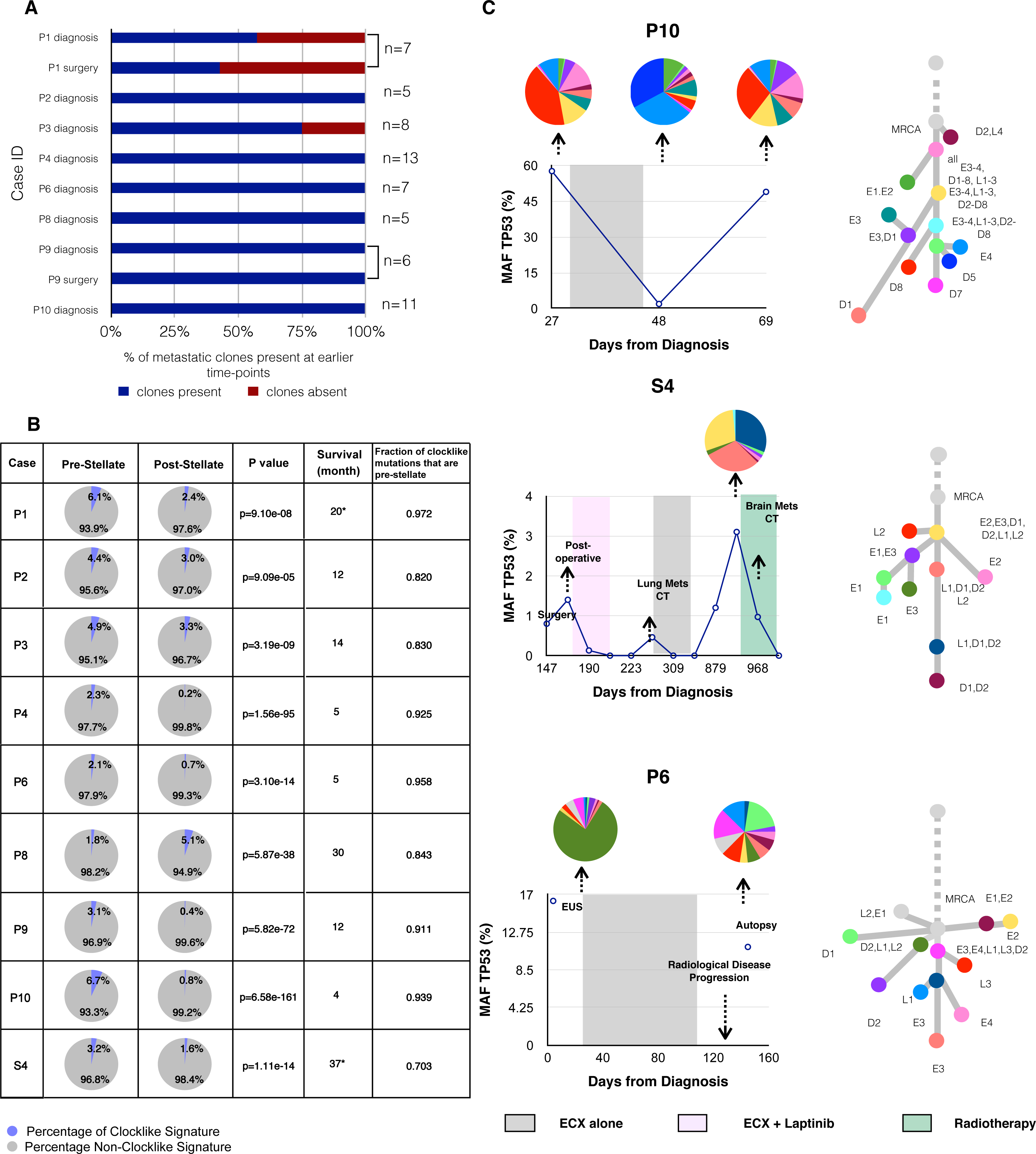
Temporal tracing of metastatic EAC using multiple lines of evidence. A) Proportion of metastatic subclones present at earlier time-points in archival formalin fixed paraffin embedded (FFPE) samples. The case ID is shown on the Y-axis along with the time-point that the sample was taken, and the % of metastatic subclones present on the X-axis. The n represents the total number of metastatic subclones. B) Mutational signature analysis of ageing signature (signature 1) pre-and post-diaspora in all 8 cases with local and distant spread (p<1.18e-90 for all cases). Chi Squared test was used to determine the p value. Survival is shown in months from the point of diagnosis *= cases which underwent surgery. C) Plasma ctDNA 1X WGS and digital droplet PCR (ddPCR) analysis for TP53 mutant allele fraction (MAF). The MAF of TP53 (%) is shown on the Y-axis and days from diagnosis are shown on the X-axis. The shaded areas represent time periods of therapy. 1X WGS at select time-points was performed and the clonal composition of these samples is shown as pie-charts. The color of each subclone corresponds to the color of the corresponding node on the adjacent phylogenetic tree.

Next we investigated eight cases (P1-4, P6, P8-10) for which the esophageal diagnostic biopsy (FFPE tissue) was available, with a median time prior to autopsy of 12 months (range 5-30 months, Figure 3A). 1X WGS identified metastatic subclones detected at autopsy in the diagnostic sample, ranging from 100% of subclones in 6 cases (P2, P4, P6, P8, P9, P10) to 75% in P3 and 57% in P1. This analysis also clarified that all metastatic subclones in P9 (Figure 1 identified from 50X WGS in the clonal discovery part of the study) arose from the primary site and were present at the earliest presentation of disease.

Tumor reseeding from metastatic sites to the primary tumor (Kim et al., 2009) or from metastasis to metastasis (Gundem et al., 2015) have been suggested as possible modes of spread. However, in this study the direction of seeding from primary to metastases, rather than vice versa, is clearly indicated by two observations. Firstly, the founder clone of the diaspora was observed in the primary tumor for all cases. If this clone originated in a metastasis, we would expect to observe additional subclones shared between multiple metastases and not the primary, but such subclones are not observed. Secondly, the founder clone of each diaspora and the great majority of subclones below it in the phylogenetic tree were identified in primary diagnostic samples, indicating that this clone was already present in each primary tumor at diagnosis. Metastasis to metastasis seeding would result in a subclone present in multiple metastases and absent from the primary tumor. When we included all 1X WGS and 50X WGS samples (8 and 4 samples, respectively, from the primary), no such subclones were identified in our cases.

In eight cases, plasma was available from rapid autopsy and 1X WGS of circulating tumor DNA (ctDNA) demonstrated that in all but one case (P1), every subclone from autopsy was also represented in plasma (Figure S8). In P1, the three subclones not found in the plasma were distal sub-clonal branches on the phylogenetic tree. We also assessed the clonal composition of ctDNA at earlier time-points in five available cases and assessed the TP53 fraction using digital PCR (Figure 3C, Figure S9, Table S12). All clones and subclones from the 50X WGS phylogenetic tree were detected at earlier time points in S4, P6, P10, (Figure 3C), while 83% and 29% of metastatic subclones were detected in S3 and P1 (Figure S9), respectively. Interestingly, P6 was a patient being treated with curative intent and had no radiological evidence of distant nodal or organ metastases at the time of clinical staging. However, at the time of diagnosis all subclones later found in the metastases were already present in the blood plasma (Figure 3C). Case S4 is noteworthy as the brain metastases (D1, D2 in Figure 1) appeared to have originated from a subclone shared between the primary and a local lymph node, both of which were removed at the time of surgery (Figure 3C).

## Discussion

We have gathered multiple lines of evidence which suggest that for the majority of EACs metastasis occurs rapidly to multiple sites. These lines of evidence can be summarized as follows. We observe multiple subclones each seeding multiple metastatic sites. These subclones are frequently derived from a single parental clone, often resulting in a stellate pattern on the phylogenetic tree. Metastases in solid organs can bypass nodal involvement. Samples within solid organ sites frequently resemble distant metastases more closely than neighboring metastases within the same organ, i.e. no tropism is observed. All metastases appear to have spread directly from the primary site, with little or no evidence of metastasis-to-metastasis seeding. One interesting possibility is that because the esophagus is highly vascularized, it may be particularly subject to hematogenous spread.

These features differ in some respects from previously described models of metastasis and we propose that they may constitute a distinct model of evolution. We suggest that this cancer (Pienta et al., 2013). Within this context, it is associated with the observation that multiple cell populations in metastatic sites are directly linked to the primary site of origin and that individual subclones seed multiple tissue types, analogous to a diaspora crossing multiple national boundaries.

A number of features were frequently associated with this phenomenon (Figure 4), with 9 of the cases (all except S3) displaying at least 2 of the 4 following features: i) stellate pattern on the phylogenetic tree; ii) lack of signature 1 mutations post MRCA or post-diaspora; iii) spread of subclones to multiple organs of different type; iv) evidence for selection in post diaspora genotypes. Regarding the latter, we looked for driver amplifications post MRCA or post diaspora on a per case basis and identified selection in 6/10 cases. However, this is likely to be an under-estimate, since there may be non-copy number drivers present in additional cases. The ratio of non-synonymous to synonymous mutations (dN/dS) analyzed across all cases as a whole, (Dentro et al., 2018), indicated positive selection in both clonal and subclonal genomes, albeit with lower levels of selection within subclones (Figure S10).

**Figure 4.**
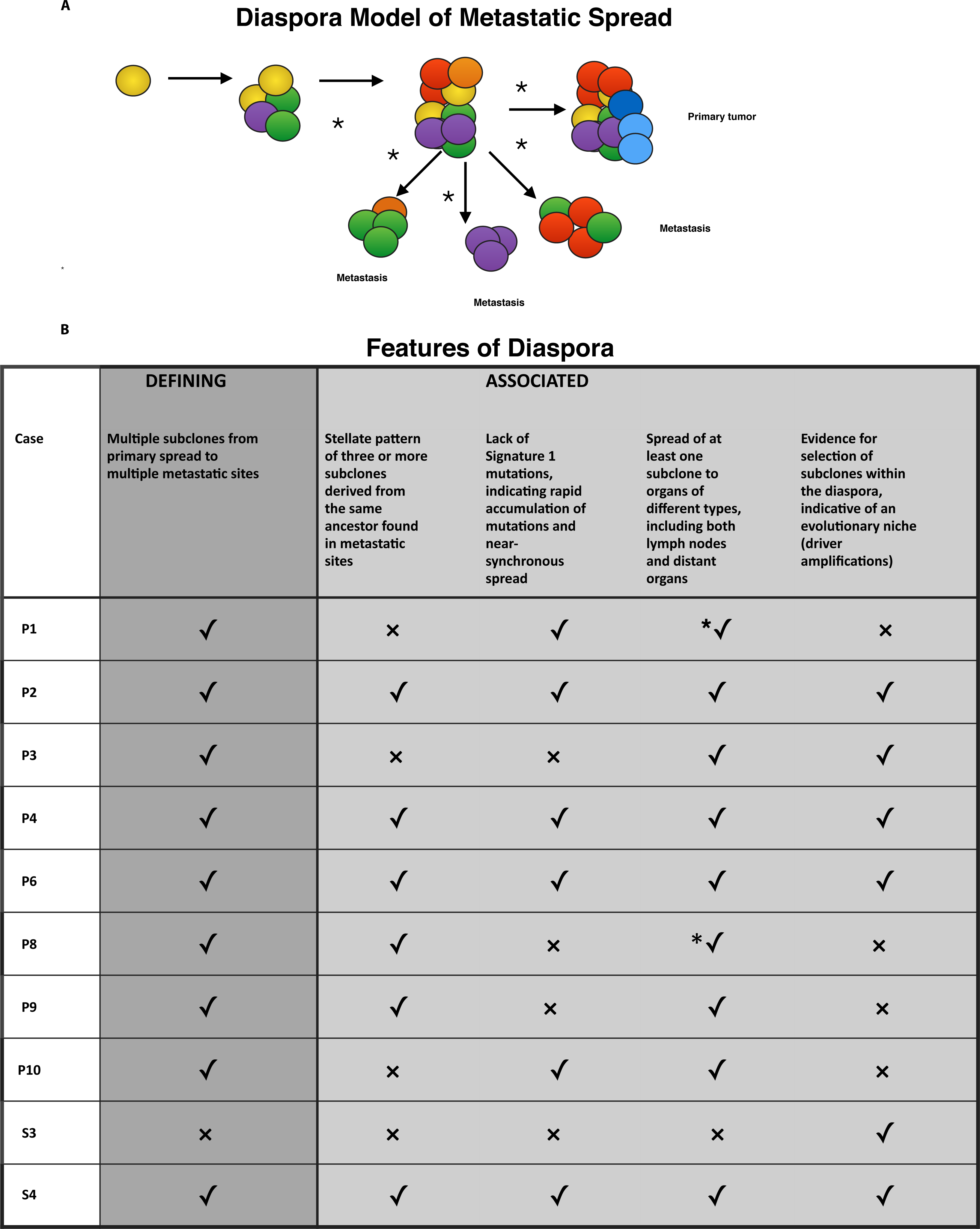
Diaspora model of metastatic spread and associated features. Panel A depicts clonal diaspora with colored circles representing clones and subclones. *= evidence of selection. Panel B explains the five features seen in diaspora (one is defining, and the other are associated with diaspora) and whether these are present (✓) or absent (x) in each case. *✓ implies that the feature is present, and that the evidence was from 1X WGS.

Until recently the genomic architectures of metastatic samples have not been defined with enough resolution to discern temporal patterns of metastatic spread. Several distinct patterns are now emerging which are not necessarily mutually exclusive or cancer-type specific. In pancreatic cancer, Yachida et al. demonstrated that distant organ seeding was a late event consistent with a linear progression model (Yachida et al., 2010). In prostate cancer, linear progression is often succeeded by multiple waves of seeding (Gundem et al., 2015). The same study further demonstrated widespread subclonal evolution in metastases and metastasis-to-metastasis spread in keeping with the relatively long longevity of prostate cancer. Strikingly, a stellate pattern was not observed in any of the cases in that study, despite using a similar design to that used in this study.

In Table S13 we compare the features of our proposed Diaspora model to the previously posited linear (Foulds, 1954) and parallel (Klein, 2009) models and consider the implications for clinical practice. Whereas the linear model predicts that linear progression will lead to a single subclone seeding lymph node sites followed by transmission to distant organs, the diaspora model posits simultaneous seeding of multiple sites directly from the primary. Unlike the parallel model, the diaspora model implies that metastasis formation occurs after the majority of evolution has occurred in the primary tumor, resulting in multiple subclones found in common between primary and metastatic tumors. Contemporaneous with this study, lymphatic and distant metastases in colon cancer have been shown to arise from independent subclones in the primary tumor with disparate evolutionary trajectories (Naxerova et al., 2017). In contrast, in EAC we find that individual subclones frequently seed both lymph node and distant organs suggesting that disparate trajectories for nodal and solid organ metastases do not exist for this disease (Figure 1, 2). Of note we acknowledge that, despite the extensive and systematic sampling across all autopsy cases, further sampling may add further branches to our phylogenetic tree although this is unlikely to affect the diaspora event itself.

In common with the Big Bang Model proposed for colorectal cancer (Sottoriva et al., 2015), our model predicts the occurrence of highly branching phylogenies. However, the Big Bang Model proposes neutral dynamics, whereas we observe strong evidence for selection in subclonal populations in the form of dN/dS ratios and the occurrence of subclonal driver amplifications (Figure 1, Figures S10 and S11). Moreover, the clonal maps of the primary tumor demonstrate subclones that occupy spatially discrete areas of the primary tumor (Figure 2), in contrast to the intermixed subclones predicted by the Big Bang Model (Sottoriva et al., 2015).

The sequence of events in metastatic progression has far-reaching clinical implications (Table S13). Clonal architecture in EAC defies anatomical location of lymph node stations and distant sites, which is the current basis for the TNM staging and determines whether curative therapy is appropriate. It has been suggested that the high recurrence rate, 52% within one year, results from seeding of distant metastases that are not detected at the time of diagnosis (Mariette et al., 2003). This study provides molecular evidence for this observation and highlights the need for different systemic approaches to disease management, including more aggressive adjuvant therapy which is not currently the mainstay of treatment (Burt et al., 2017; Gabriel et al., 2017; Pasquali et al., 2017; Sjoquist et al., 2011). Furthermore, the presence of subclones identified from autopsy in diagnostic blood and tissue samples suggests that there is the potential for earlier detection.

The occurrence of metastasis is a pivotal event in the life history of a cancer. Understanding the mechanism behind such an event would have potential relevance to predicting and preventing metastatic spread. From the relatively modest number of cases within this study, we were not able to identify aberrations in specific genes or recurrent copy number changes associated with the occurrence of metastasis (Schumacher et al., 2017; Stoecklein et al., 2008). In a number of cases, diaspora was coincident with an increase in the proportion of signature 3 mutations, associated with failure of DNA double-strand break-repair by homologous recombination. Our findings are in keeping with the failure of DNA repair driving the appearance of genomic heterogeneity. Whether the heterogeneity observed is itself the driver of diaspora or merely a symptom is an important area for future study. Our investigations of the potential drivers of diaspora were limited to genomic factors, and further multi-platform studies looking at epigenetic and transcriptomic factors are other important avenues of future research. We anticipate that analyses of single cells or small clusters from primary sites, disseminated tumor cells and circulating tumor cells will also yield finer resolution of the processes of dissemination and metastasis. In addition, understanding the timing of this event will be of value in planning key events in patient management such as surgery and oncological therapy.

In cancer generally there are currently very few in-depth studies examining the spatial and temporal evolution of metastases as noted in a recent comprehensive study of metastases from multiple primary sites (Robinson et al., 2017). Further studies are required to ascertain whether our diaspora theory also pertains to other cancer types.

## Acknowledgements

We would like to thank the following individuals for their help with study set-up, patient liaison and tissue collection, Ben Smith, Nyrai Chinyama, Vijay Sujendran, Peter Safranek, Athanosios Xanthos, Tara Nuckcheddy-Grant, Rachel de la Rue, Sebastian Zeki, Rachael Fels Elliott, Peter Collins, Kitty Puttock, Sophie Rabey and staff at Arthur Rank Hospice and Luke A Wylie for scientific discussion and contribution. We are grateful to Professor Simon Tavaré, FRS for his guidance and support for the esophageal whole genome sequencing thank Jo Westmoreland, LMB visual aids for her graphic art expertise. Thanks also go to the Cancer Research UK Cambridge Institute Genomics Core for their technical expertise. Above all, we are indebted to the patients who donated tissue samples to this project, and their families who supported them through it.

## Author Contributions

AN designed the study, implemented the rapid autopsy study, performed the experiments, analyzed data and wrote the manuscript. MG and S.D.P contributed expertise in pathology and sample collection for the rapid autopsy study. ID-B and NG assisted in study implementation, and along with JC, assisted with sample collection at autopsy. M.D.E performed genomic data generation and QC. LB conducted data management. XL, PL-S and JW were involved with autopsy sample collection, advice on experiments and data analysis, and XL contributed to paper writing. LA and IM assisted with data analysis. NG assisted with study Implementation. SMac coordinated the sequencing of samples from the OCCAMS project and contributed to paper writing. SM and AM provided pathology data. TT, SG, LP and DG assisted in implementation and ethical conduct of the autopsy study. R.H.H and AH were involved in surgical sample collection and providing surgical expertise. M.R.S contributed to critical evaluation of the study data and manuscript. D.C.W was responsible for data analysis, paper writing, and assuring integrity of data. The OCCAMS consortium was the vehicle through which the infrastructure and funding was obtained to support the study and the consortium contributed to discussions on the ICGC data and the clinical ramifications. R.C.F provided grant funding and was responsible for study design, supervision of the project, writing the paper and assuring integrity of the data.

The authors declare no competing interests.

## Methods

### Patients and tumor samples

We collected 396 samples from surgery and endoscopy (part of esophageal ICGC) as well as from a rapid autopsy programme called PHOENIX. Patients were eligible if they were at least 18 years of age and had received a confirmed diagnosis of EAC following central pathology review. Patients were only approached for the PHOENIX study following a palliative diagnosis, with the full involvement of the multidisciplinary team. All demographic and clinical data was anonymized and stored on a central study database (OpenClinica and Labkey).

All samples were collected according to a strict SOP. Post-mortems were completed within 6 hours of death to ensure tissue integrity for WGS. The clinical characteristics of the patients are provided in Tables S2 and S3.

### Whole genome sequencing and data analysis

We used the Illumina HiSeq platform to perform WGS on multiple regions collected from each primary tumor, lymph node and/or solid organ metastasis (Figure S1A, B, Tables S4 and S5). All DNA extractions and WGS conformed with ICGC quality control standards and required ≥70% cellularity and a matched germline sample. WGS was performed at high depth (median coverage 66.3, IQR 56.1-87.2) to discover mutations in 122 samples from 18 patients (Tables S3a and b). In addition, low depth WGS (median coverage 1, IQR 1-5) was performed to track these mutations spatially in up to 48 solid tissue samples per case, (total=250) and 8 ctDNA samples at autopsy. Temporal tracking was performed in cases with archival biopsy material, and where historical bloods were available (Table S12, Figure 3A, C). For each patient the number of subclones and the cancer cell fraction within each subclone was inferred using an extension of a previously described Bayesian Dirichlet process (Nik-Zainal et al., 2012) and we applied a set of previously described rules to derive a phylogenetic tree (Additional Methods; (Jiao et al., 2014). All sequencing data have been deposited in the European Genome-Phenome Archive under accession number EGAD00001003403. TP53 analysis in cell free tumor DNA (ctDNA) was performed using Digital PCR on the Bio-rad platform (Bio-rad, California) using validated TP53 assays (Table S14).

## References

Ajani, J.A., D’Amico, T.A., Almhanna, K., Bentrem, D.J., Besh, S., Chao, J., Das, P., Denlinger, C., Fanta, P., Fuchs, C.S., et al. (2015). Esophageal and esophagogastric junction cancers, version 1.2015. J Natl Compr Canc Netw 13, 194-227.

Alexandrov, L.B., Jones, P.H., Wedge, D.C., Sale, J.E., Campbell, P.J., Nik-Zainal, S., and Stratton, M.R. (2015). Clock-like mutational processes in human somatic cells. Nat Genet 47, 1402-1407.

Alexandrov, L.B., Nik-Zainal, S., Wedge, D.C., Aparicio, S.A., Behjati, S., Biankin, A.V., Bignell, G.R., Bolli, N., Borg, A., Borresen-Dale, A.L., et al. (2013). Signatures of mutational processes in human cancer. Nature 500, 415-421.

Behjati, S., Gundem, G., Wedge, D.C., Roberts, N.D., Tarpey, P.S., Cooke, S.L., Van Loo, P., Alexandrov, L.B., Ramakrishna, M., Davies, H., et al. (2016). Mutational signatures of ionizing radiation in second malignancies. Nat Commun 7, 12605.

Blokzijl, F., de Ligt, J., Jager, M., Sasselli, V., Roerink, S., Sasaki, N., Huch, M., Boymans, S., Kuijk, E., Prins, P., et al. (2016). Tissue-specific mutation accumulation in human adult stem cells during life. Nature 538, 260-264.

Bolli, N., Avet-Loiseau, H., Wedge, D.C., Van Loo, P., Alexandrov, L.B., Martincorena, I., Dawson, K.J., Iorio, F., Nik-Zainal, S., Bignell, G.R., et al. (2014). Heterogeneity of genomic evolution and mutational profiles in multiple myeloma. Nat Commun 5, 2997.

Burt, B.M., Groth, S.S., Sada, Y.H., Farjah, F., Cornwell, L., Sugarbaker, D.J., and Massarweh, N.N. (2017). Utility of Adjuvant Chemotherapy After Neoadjuvant Chemoradiation and Esophagectomy for Esophageal Cancer. Ann Surg 266, 297-304.

Chen, X., Schulz-Trieglaff, O., Shaw, R., Barnes, B., Schlesinger, F., Kallberg, M., Cox, A.J., Kruglyak, S., and Saunders, C.T. (2016). Manta: rapid detection of structural variants and indels for germline and cancer sequencing applications. Bioinformatics 32, 1220-1222.

Cunningham, D., Starling, N., Rao, S., Iveson, T., Nicolson, M., Coxon, F., Middleton, G., Daniel, F., Oates, J., Norman, A.R., et al. (2008). Capecitabine and oxaliplatin for advanced esophagogastric cancer. N Engl J Med 358, 36-46.

Davis, A., Gao, R., and Navin, N. (2017). Tumor evolution: Linear, branching, neutral or punctuated? Biochim Biophys Acta 1867, 151-161.

Dentro, S.C., Leshchiner, I., Haase, K., Tarabichi, M., Wintersinger, J., Deshwar, A.G., Yu, K., Rubanova, Y., Macintyre, G., Vazquez-Garcia, I., et al. (2018). Portraits of genetic intra-tumour heterogeneity and subclonal selection across cancer types. bioRxiv.

Dulak, A.M., Schumacher, S.E., van Lieshout, J., Imamura, Y., Fox, C., Shim, B., Ramos, A.H., Saksena, G., Baca, S.C., Baselga, J., et al. (2012). Gastrointestinal adenocarcinomas of the esophagus, stomach, and colon exhibit distinct patterns of genome instability and oncogenesis. Cancer Res 72, 4383-4393.

Dulak, A.M., Stojanov, P., Peng, S., Lawrence, M.S., Fox, C., Stewart, C., Bandla, S., Imamura, Y., Schumacher, S.E., Shefler, E., et al. (2013). Exome and whole-genome sequencing of esophageal adenocarcinoma identifies recurrent driver events and mutational complexity. Nat Genet 45, 478-486.

Foulds, L. (1954). The experimental study of tumor progression: a review. Cancer Res 14, 327-339.

Frankell, A.M., Jammula, S., Contino, G., Killcoyne, S.S., Abbas, S., Perner, J., Bower, L., Devonshire, G., Grehan, N., Mok, J., et al. (2018). The landscape of selection in 551 Esophageal Adenocarcinomas defines genomic biomarkers for the clinic. bioRxiv.

Gabriel, E., Attwood, K., Shah, R., Nurkin, S., Hochwald, S., and Kukar, M. (2017). Novel Calculator to Estimate Overall Survival Benefit from Neoadjuvant Chemoradiation in Patients with Esophageal Adenocarcinoma. J Am Coll Surg 224, 884-894 e881.

Gao, Z., Wyman, M.J., Sella, G., and Przeworski, M. (2016). Interpreting the Dependence of Mutation Rates on Age and Time. PLoS Biol 14, e1002355.

Genomes Project, C., Abecasis, G.R., Auton, A., Brooks, L.D., DePristo, M.A., Durbin, R.M., Handsaker, R.E., Kang, H.M., Marth, G.T., and McVean, G.A. (2012). An integrated map of genetic variation from 1,092 human genomes. Nature 491, 56-65.

Gerstung, M., Jolly, C., Leshchiner, I., Dentro, S.C., Gonzalez, S., Mitchell, T.J., Rubanova, Y., Anur, P., Rosebrock, D., Yu, K., et al. (2017). The evolutionary history of 2,658 cancers. bioRxiv.

Greaves, M., and Maley, C.C. (2012). Clonal evolution in cancer. Nature 481, 306-313.

Gundem, G., Van Loo, P., Kremeyer, B., Alexandrov, L.B., Tubio, J.M.C., Papaemmanuil, E., Brewer, D.S., Kallio, H.M.L., Hognas, G., Annala, M., et al. (2015). The evolutionary history of lethal metastatic prostate cancer. Nature 520, 353-357.

Japanese Gastric Cancer, A. (2011). Japanese classification of gastric carcinoma: 3rd English edition. Gastric Cancer 14, 101-112.

Jiao, W., Vembu, S., Deshwar, A.G., Stein, L., and Morris, Q. (2014). Inferring clonal evolution of tumors from single nucleotide somatic mutations. BMC Bioinformatics 15, 35.

Kim, M.Y., Oskarsson, T., Acharyya, S., Nguyen, D.X., Zhang, X.H., Norton, L., and Massague, J. (2009). Tumor self-seeding by circulating cancer cells. Cell 139, 1315-1326.

Klein, C.A. (2009). Parallel progression of primary tumours and metastases. Nat Rev Cancer 9, 302-312.

Kuipers, J., Jahn, K., Raphael, B.J., and Beerenwinkel, N. (2017). Single-cell sequencing data reveal widespread recurrence and loss of mutational hits in the life histories of tumors. Genome Res 27, 1885-1894.

Letouze, E., Shinde, J., Renault, V., Couchy, G., Blanc, J.F., Tubacher, E., Bayard, Q., Bacq, D., Meyer, V., Semhoun, J., et al. (2017). Mutational signatures reveal the dynamic interplay of risk factors and cellular processes during liver tumorigenesis. Nat Commun 8, 1315.

Li, H., and Durbin, R. (2009). Fast and accurate short read alignment with Burrows-Wheeler transform. Bioinformatics 25, 1754-1760.

Liu, D., Abbosh, P., Keliher, D., Reardon, B., Miao, D., Mouw, K., Weiner-Taylor, A., Wankowicz, S., Han, G., Teo, M.Y., et al. (2017). Mutational patterns in chemotherapy resistant muscle-invasive bladder cancer. Nat Commun 8, 2193.

Lodato, M.A., Rodin, R.E., Bohrson, C.L., Coulter, M.E., Barton, A.R., Kwon, M., Sherman, M.A., Vitzthum, C.M., Luquette, L.J., Yandava, C.N., et al. (2018). Aging and neurodegeneration are associated with increased mutations in single human neurons. Science 359, 555-559.

Lou, F., Sima, C.S., Adusumilli, P.S., Bains, M.S., Sarkaria, I.S., Rusch, V.W., and Rizk, N.P. (2013). Esophageal cancer recurrence patterns and implications for surveillance. J Thorac Oncol 8, 1558-1562.

Mariette, C., Balon, J.M., Piessen, G., Fabre, S., Van Seuningen, I., and Triboulet, J.P. (2003). Pattern of recurrence following complete resection of esophageal carcinoma and factors predictive of recurrent disease. Cancer 97, 1616-1623.

Martincorena Inigo, R.K.M., Gerstung Moritz, Dawson Kevin J, Haase Kerstin, Van Loo Peter, Davies Helen, Michael R. Stratton Michael R, Campbell Peter J. (2017). Universal Patterns Of Selection In Cancer And Somatic Tissues. Cell.

Matsuda, S., Takeuchi, H., Kawakubo, H., and Kitagawa, Y. (2017). Three-field lymph node dissection in esophageal cancer surgery. J Thorac Dis 9, S731-S740.

Murugaesu, N., Wilson, G.A., Birkbak, N.J., Watkins, T., McGranahan, N., Kumar, S., Abbassi-Ghadi, N., Salm, M., Mitter, R., Horswell, S., et al. (2015). Tracking the genomic evolution of esophageal adenocarcinoma through neoadjuvant chemotherapy. Cancer Discov 5, 821-831.

Naxerova, K., Reiter, J.G., Brachtel, E., Lennerz, J.K., van de Wetering, M., Rowan, A., Cai, T., Clevers, H., Swanton, C., Nowak, M.A., et al. (2017). Origins of lymphatic and distant metastases in human colorectal cancer. Science 357, 55-60.

Nik-Zainal, S., Van Loo, P., Wedge, D.C., Alexandrov, L.B., Greenman, C.D., Lau, K.W., Raine, K., Jones, D., Marshall, J., Ramakrishna, M., et al. (2012). The life history of 21 breast cancers. Cell 149, 994-1007.

Nones, K., Waddell, N., Wayte, N., Patch, A.M., Bailey, P., Newell, F., Holmes, O., Fink, J.L., Quinn, M.C., Tang, Y.H., et al. (2014). Genomic catastrophes frequently arise in esophageal adenocarcinoma and drive tumorigenesis. Nat Commun 5, 5224.

Nowell, P.C. (1976). The clonal evolution of tumor cell populations. Science 194, 23-28.

Pasquali, S., Yim, G., Vohra, R.S., Mocellin, S., Nyanhongo, D., Marriott, P., Geh, J.I., and Griffiths, E.A. (2017). Survival After Neoadjuvant and Adjuvant Treatments Compared to Surgery Alone for Resectable Esophageal Carcinoma: A Network Meta-analysis. Ann Surg 265, 481-491.

Pienta, K.J., Robertson, B.A., Coffey, D.S., and Taichman, R.S. (2013). The cancer diaspora: Metastasis beyond the seed and soil hypothesis. Clin Cancer Res 19, 5849-5855.

Robinson, D.R., Wu, Y.M., Lonigro, R.J., Vats, P., Cobain, E., Everett, J., Cao, X., Rabban, E., Kumar-Sinha, C., Raymond, V., et al. (2017). Integrative clinical genomics of metastatic cancer. Nature 548, 297-303.

Ross-Innes, C.S., Becq, J., Warren, A., Cheetham, R.K., Northen, H., O’Donovan, M., Malhotra, S., di Pietro, M., Ivakhno, S., He, M., et al. (2015). Whole-genome sequencing provides new insights into the clonal architecture of Barrett’s esophagus and esophageal adenocarcinoma. Nat Genet 47, 1038-1046.

Saunders, C.T., Wong, W.S., Swamy, S., Becq, J., Murray, L.J., and Cheetham, R.K. (2012). Strelka: accurate somatic small-variant calling from sequenced tumor-normal sample pairs. Bioinformatics 28, 1811-1817.

Scheinin, I., Sie, D., Bengtsson, H., van de Wiel, M.A., Olshen, A.B., van Thuijl, H.F., van Essen, H.F., Eijk, P.P., Rustenburg, F., Meijer, G.A., et al. (2014). DNA copy number analysis of fresh and formalin-fixed specimens by shallow whole-genome sequencing with identification and exclusion of problematic regions in the genome assembly. Genome Res 24, 2022-2032.

Schumacher, S., Bartenhagen, C., Hoffmann, M., Will, D., Fischer, J.C., Baldus, S.E., Vay, C., Fluegen, G., Dizdar, L., Vallbohmer, D., et al. (2017). Disseminated tumour cells with highly aberrant genomes are linked to poor prognosis in operable oesophageal adenocarcinoma. Br J Cancer 117, 725-733.

Secrier, M., Li, X., de Silva, N., Eldridge, M.D., Contino, G., Bornschein, J., MacRae, S., Grehan, N., O’Donovan, M., Miremadi, A., et al. (2016). Mutational signatures in esophageal adenocarcinoma define etiologically distinct subgroups with therapeutic relevance. Nat Genet 48, 1131-1141.

Sjoquist, K.M., Burmeister, B.H., Smithers, B.M., Zalcberg, J.R., Simes, R.J., Barbour, A., Gebski, V., and Australasian Gastro-Intestinal Trials, G. (2011). Survival after neoadjuvant chemotherapy or chemoradiotherapy for resectable oesophageal carcinoma: an updated meta-analysis. Lancet Oncol 12, 681-692.

Smyth, E.C., Lagergren, J., Fitzgerald, R.C., Lordick, F., Shah, M.A., Lagergren, P., and Cunningham, D. (2017). Oesophageal cancer. Nat Rev Dis Primers 3, 17048.

Sottoriva, A., Kang, H., Ma, Z., Graham, T.A., Salomon, M.P., Zhao, J., Marjoram, P., Siegmund, K., Press, M.F., Shibata, D., et al. (2015). A Big Bang model of human colorectal tumor growth. Nat Genet 47, 209-216.

Sottoriva, A., Spiteri, I., Piccirillo, S.G., Touloumis, A., Collins, V.P., Marioni, J.C., Curtis, C., Watts, C., and Tavare, S. (2013). Intratumor heterogeneity in human glioblastoma reflects cancer evolutionary dynamics. Proc Natl Acad Sci U S A 110, 4009-4014.

Sporn, M.B. (1996). The war on cancer. Lancet 347, 1377-1381.

Stiles, B.M., Nasar, A., Mirza, F.A., Lee, P.C., Paul, S., Port, J.L., and Altorki, N.K. (2012). Worldwide Oesophageal Cancer Collaboration guidelines for lymphadenectomy predict survival following neoadjuvant therapy. Eur J Cardiothorac Surg 42, 659-664.

Stoecklein, N.H., Hosch, S.B., Bezler, M., Stern, F., Hartmann, C.H., Vay, C., Siegmund, A., Scheunemann, P., Schurr, P., Knoefel, W.T., et al. (2008). Direct genetic analysis of single disseminated cancer cells for prediction of outcome and therapy selection in esophageal cancer. Cancer Cell 13, 441-453.

Waterman, T.A., Hagen, J.A., Peters, J.H., DeMeester, S.R., Taylor, C.R., and Demeester, T.R. (2004). The prognostic importance of immunohistochemically detected node metastases in resected esophageal adenocarcinoma. Ann Thorac Surg 78, 1161-1169; discussion 1161-1169.

Weaver, J.M., Ross-Innes, C.S., Shannon, N., Lynch, A.G., Forshew, T., Barbera, M., Murtaza, M., Ong, C.A., Lao-Sirieix, P., Dunning, M.J., et al. (2014). Ordering of mutations in preinvasive disease stages of esophageal carcinogenesis. Nat Genet 46, 837-843.

Yachida, S., Jones, S., Bozic, I., Antal, T., Leary, R., Fu, B., Kamiyama, M., Hruban, R.H., Eshleman, J.R., Nowak, M.A., et al. (2010). Distant metastasis occurs late during the genetic evolution of pancreatic cancer. Nature 467, 1114-1117.

Yates, L.R., Gerstung, M., Knappskog, S., Desmedt, C., Gundem, G., Van Loo, P., Aas, T., Alexandrov, L.B., Larsimont, D., Davies, H., et al. (2015). Subclonal diversification of primary breast cancer revealed by multiregion sequencing. Nat Med 21, 751-759.

